# Systems biology analysis identifies TCF7L1 as a key regulator of metastasis in Ewing sarcoma

**DOI:** 10.1101/2021.02.25.432862

**Authors:** Florencia Cidre-Aranaz, Jing Li, Tilman L. B. Hölting, Martin F. Orth, Roland Imle, Stefanie Kutschmann, Giulia Ammirati, Katharina Ceranski, Martha Julia Carreño-Gonzalez, Merve Kasan, Aruna Marchetto, Cornelius M. Funk, Felix Bestvater, Simone Bersini, Chiara Arrigoni, Matteo Moretti, Laura Romero-Pérez, Ana Banito, Shunya Ohmura, Julian Musa, Thomas Kirchner, Maximilian M. L. Knott, Thomas G. P. Grünewald

**Affiliations:** Hopp-Children’s Cancer Center (KiTZ), Heidelberg, Germany; Division of Translational Pediatric Sarcoma Research (B410), German Cancer Research Center (DKFZ), German Cancer Consortium (DKTK), Heidelberg, Germany; Max-Eder Research Group for Pediatric Sarcoma Biology, Institute of Pathology, Faculty of Medicine, LMU Munich, Munich, Germany; Soft-Tissue Sarcoma Junior Research Group, German Cancer Research Center (DKFZ), German Cancer Consortium (DKTK), Heidelberg, Germany; Faculty of Biosciences, Heidelberg University, Heidelberg, Germany; Division of Pediatric Surgery, Department of General, Visceral and Transplantation Surgery, University Hospital Heidelberg, Heidelberg, Germany; Regenerative Medicine Technologies Laboratory, Ente Ospedaliero Cantonale (EOC), Lugano, Switzerland; Light Microscopy Facility (W210), German Cancer Research Center (DKFZ), German Cancer Consortium (DKTK), Heidelberg, Germany; Biomedical Sciences Faculty, Università della Svizzera Italiana (USI), Lugano, Switzerland; Department of General, Visceral and Transplantation Surgery, Heidelberg University Hospital, Heidelberg, Germany; Institute of Pathology, Heidelberg University Hospital, Heidelberg, Germany

**Keywords:** Ewing sarcoma, TCF7L1, pediatric oncology, metastasis, systems biology

## Abstract

Identification of cancer stemness genes is crucial to understanding the underlying biology of therapy resistance, relapse, and metastasis. Ewing sarcoma (EwS) is the second most common bone tumor in children and adolescents. It is a highly aggressive cancer associated with a dismal survival rate (<30%) for patients with metastatic disease at diagnosis (∼25% of cases). Hence, deciphering the underlying mechanisms of metastasis is imperative. EwS tumors are characterized by a remarkably ‘silent’ genome with a single driver mutation generating an oncogenic fusion transcription factor (*EWSR1-ETS*). Thus, EwS constitutes an ideal model to study how perturbation of a transcriptional network by a dominant oncogene can mediate metastasis, even though canonical metastasis-associated genes are not mutated.

Here, through the implementation of an integrative systems biology approach, we identified transcription factor 7 like 1 (*TCF7L1*, alias *TCF3*) as a prognostically-relevant and *EWSR1-ETS* suppressed determinant of metastasis in EwS. We demonstrated that conditional *TCF7L1* re-expression significantly reduces EwS single-cell migration, invasion and anchorage-independent growth in 3D assays *in vitro*, and tumorigenesis *in vivo* mediated by its DNA binding domain. In primary EwS tumors as well as in functional orthotopic *in vivo* models, low *TCF7L1* expression was associated with pro-metastatic gene signatures and a much higher migratory and metastatic capacity of EwS cells, which correlated with poor outcome of EwS patients.

Collectively, our findings establish TCF7L1 as a major regulator of metastasis in EwS, which may be utilized as a prognostic biomarker and open inroads to future therapeutic intervention.

## INTRODUCTION

Cancer stemness genes are a cornerstone governing therapy resistance, relapse, and metastasis^1,2^. In carcinomas, many stemness/metastasis genes and associated pathways are frequently activated through somatic mutations including copy-number changes and base substitutions^3,4^. Surprisingly, pediatric tumors –despite being largely composed of undifferentiated stem cell-like cells– typically present low numbers of mutations that generally do not involve canonical stemness/metastasis-associated genes^5^.

Ewing sarcoma (EwS) is a highly aggressive bone and soft-tissue cancer with great propensity for early hematological metastasis mainly affecting children, adolescents, and young adults^6^. In past decades, the development and implementation of multimodal chemotherapeutic regimens for the treatment of patients with localized (i.e. non-metastatic) EwS has drastically increased overall survival^6^. However, the clinical outcome for patients with metastatic disease at diagnosis (∼ 25% of cases) or relapse remains unacceptably poor (<30% survival rates), even with the application of highly toxic therapies^7^. Hence, it is crucial to unravel the underlying biologic mechanisms that drive stemness and metastasis in EwS to open new therapeutic avenues.

Genetically, EwS ranges among the tumors with the fewest mutations identified to date^8^. In fact, most EwS tumors exhibit only a single recurrent mutation, that is a chromosomal translocation fusing the *EWSR1* gene and variable members of the *ETS* family of transcription factors (TFs)^9,10^. *EWSR1-ETS* encode aberrant TFs, which massively rewire the EwS transcriptome and promote stem cell features^11–13^; even though classical stemness-and metastasis-associated genes remain not mutated^14–16^. Although EwS is genetically well characterized, its cell of origin remains unknown. Morphologically, EwS tumors resemble blastemal tissues composed of highly undifferentiated cells without overt cellular hierarchy, wherefore it is often described as an ‘embryonal tumor’^6^. Indeed, ectopic EWSR1-FLI1 expression can transcriptionally ‘reprogram’ human pediatric mesenchymal stem cells by upregulation of embryonic stem cell genes generating cells with cancer stem cell-like properties^13^.

The processes of metastasis and stemness are intimately linked^17^. In fact, accumulating evidence suggests that tumor cells of highly undifferentiated malignancies, such as EwS, may reside in a ‘metastable’ state equipping them with proliferative and migratory traits (for review see ^18^), which has been previously framed by the integrative ‘migratory stem cell’ concept^19^. Given the rather simple genetic make-up of EwS, this disease may constitute an ideal model to study how perturbation of a transcriptional network by a dominant oncogene can mediate stemness and metastasis features. Although several groups identified particular EWSR1-ETS-driven genes or pathways that likely play a role in maintenance of stemness, and hence in the establishment of a more metastatic phenotype^20^, a systematic understanding of this process and of its clinical implications is still lacking.

In the current study, we explored how EWSR1-ETS rewires the EwS transcriptome on a systems biology level, and identified the transcription factor 7 like 1 (*TCF7L1*, alias *TCF3*), a stem-cell differentiation-related transcription factor involved in the Wnt pathway^21^, as key downstream regulator of the EWSR1-ETS-mediated stemness and metastasis features. By combining preclinical and clinical data, we show that EWSR1-ETS suppresses *TCF7L1*, which promotes anchorage-independent growth and metastasis through deregulation of developmental and migration-associated gene signatures. Moreover, we show that *TCF7L1* expression is negatively correlated with poor patient outcome, collectively shedding new light on the underlying biological mechanisms governing stemness and metastasis in EwS, which may open inroads to future clinical exploitation.

## RESULTS

### Application of a systems biology approach identifies TCF7L1 as a prognostically relevant EWSR1-ETS-regulated network hub

To identify highly relevant *EWSR1-ETS* target genes involved in the stemness/metastasis axis we analyzed transcriptome profiles of 5 EwS cell lines with or without shRNA-mediated knockdown of *EWSR1-FLI1* or *-ERG* (knockdown <20% of baseline expression) for 96h according to the workflow depicted in **Fig. 1a**. This analysis yielded a list of 348 differentially expressed genes (DEGs) being up-or downregulated (|log2 FC|≥1.5) after knockdown of *EWSR1-FLI1* or *-ERG* across all cell lines (**Supplementary Table 1**). We next filtered these DEGs for genes being annotated with the gene ontology (GO) term ‘Regulation of Cell Differentiation’ using PantherDB, which was significantly enriched among the *EWSR1-ETS* regulated DEGs (*P*=4.93×10^−12^, false-discovery rate (FDR)=1.97×10^−9^). Based on the resulting 76 DEGs (**Supplementary Table 2**), we carried out a network analysis using *Cytoscape* and the *GeneMania* plugin using available information on pathway, physical, and genetic interactions^22^ (**Fig. 1b,c**). To identify key hubs within this network that may have clinical relevance and may be important for understanding downstream effects of *EWRS1-ETS*, we focused on highly interconnected TFs (*n*=11) whose expression correlated with overall survival of EwS patients. To this end, we further filtered the list of TFs for those that exhibited at least 10 interactions and that showed a significant (*P*<0.05, Mantel-Haenszel test) association with overall survival in a cohort of 166 EwS patients with matched gene expression and clinical data^23^. This analytical process yielded two TFs whose expression nominally correlated with patient overall survival (**Supplementary Table 3**). However, after adjustment for multiple testing, transcription factor 7 like 1 (*TCF7L1*, alias *TCF3*) emerged as the most promising candidate for functional follow-up (nominal *P*=0.0057; *P*=0.0228 Bonferroni-adjusted) (**Fig. 1d**). Notably, *TCF7L1* appeared to be downregulated by *EWSR1-ETS* since microarray data of each respective fusion oncogene knockdown for 96h induced on average a ∼3-fold upregulation of *TCF7L1* (**Supplementary Table 4**), which was confirmed in 5 EwS cell lines with independent assays by qRT-PCR (on average ∼6-fold upregulation (**Supplementary Fig. 1a**.). This finding was validated in xenografts of A673 cells with/without conditional knockdown of EWSR1-FLI1 (A673/TR/shEF1 cells) at the mRNA and protein level (**Supplementary Fig. 1b**).

**Fig 1.**
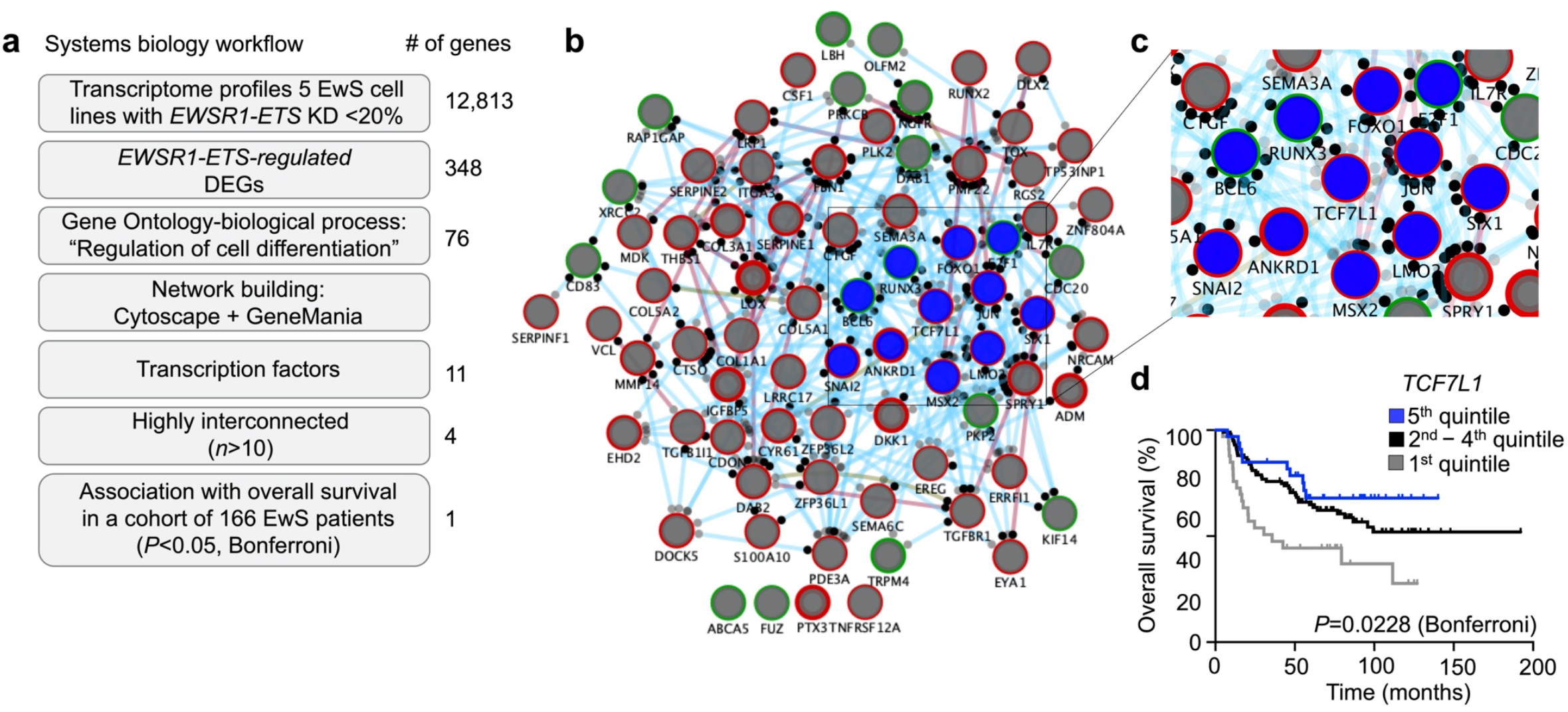
Application of a systems biology approach identifies TCF7L1 as a prognostically relevant EWSR1-ETS-regulated network hub. **a)** Workflow depicting a step-wise systems biology approach to identify DEGs regulated by EWSR1-ETS, involved in regulation of cell differentiation, functioning as highly interconnected TFs, and associated with overall survival in a cohort of 166 EwS patients. Number of genes represent remaining candidates after each filtering step. **b)** Network of EWSR1-ETS-regulated genes involved in regulation of cell differentiation. Genes are depicted as nodes (circles). Blue nodes represent TFs, node outline color show up-(green) or down-(red) regulation by EWSR1-ETS. Node border width represent strength of regulation by EWSR1-ETS (the thicker the border the higher the fold-change). Black dots surrounding the nodes represent interconnections with other nodes within the network. Connecting lines show three types of interconnection: physical (red), pathway (blue), genetic (brown). Line width represents strength of interconnection. **c)** Close-up image of highly interconnected TFs located at the center of the network (blue nodes). **d)** Kaplan-Meier survival analysis of 166 primary EwS patients stratified by quintile *TCF7L1* expression. Mantel-Haenszel test, Bonferroni corrected. DEG: differentially expressed genes; KD: knockdown.

Collectively, these data highlight *TCF7L1* as a prognostically relevant, EWSR1-ETS-regulated network hub involved in the regulation on stemness/metastasis in EwS.

### TCF7L1 re-expression inhibits tumorigenesis *in vitro* and *in vivo*

Little is known about the precise cellular function of *TCF7L1* in cancer. However, prior reports suggested that it may have either oncogenic or tumor suppressor properties depending on the cellular context^24–28^. To explore its potential role in EwS, we first analyzed the *TCF7L1* expression pattern across 18 cancer entities using publicly available microarray data from the Cancer Cell Line Encyclopedia (CCLE)^29^ and from an own study that compiled well-curated microarray data from the same cancer entities^30^. Surprisingly, *TCF7L1* was very highly, but still variably, expressed in EwS cell lines and primary tumors (**Fig. 2a**).

**Fig 2.**
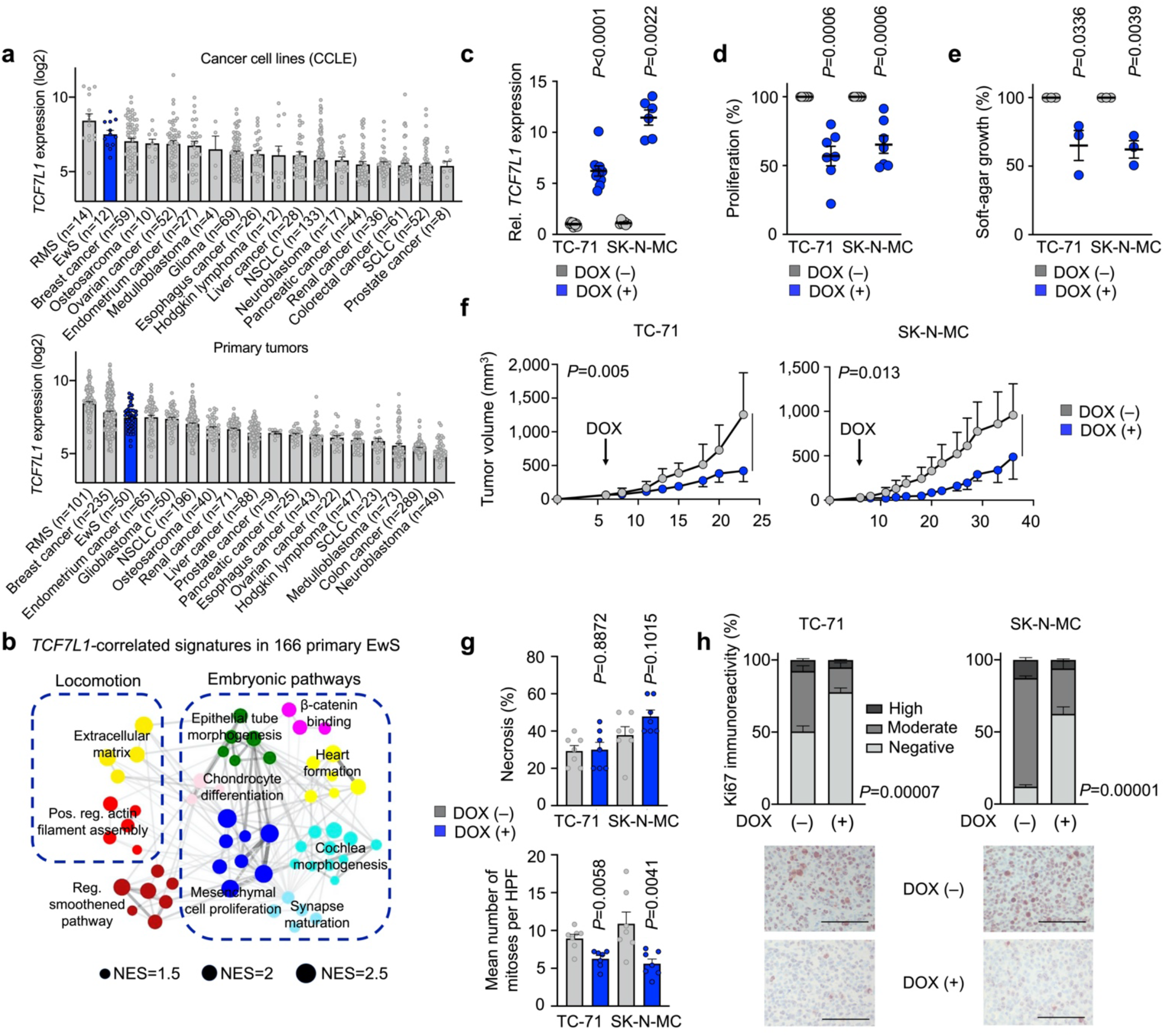
TCF7L1 re-expression inhibits tumorigenesis *in vitro* and *in vivo*. **a)** *TCF7L1* expression levels of EwS and 17 additional tumor entities in cell lines (Cancer Cell Line Encyclopedia, CCLE) or primary tumors. Data are represented as bar plots where horizontal bars represent mean and SEM. The number of samples per group (*n*) is given in parentheses. RMS, rhabdomyosarcoma; NSCLC, non-small cell lung carcinoma; SCLC: small-cell lung carcinoma. **b)** Weighted Gene Correlation Network Analysis (WGCNA) of enriched gene-sets obtained by Pearson correlation analysis of genes whose expression is negatively correlated with *TCF7L1* expression in Affymetrix gene expression data of 166 primary EwS tumors. Network depicts signatures presenting NES>1.5 and *P*<0,05. NES, normalized enrichment score. **c)** Relative *TCF7L1* expression as measured by qRT-PCR of TC-71 and SK-N-MC cells containing a DOX-inducible re-expression construct for *TCF7L1*. Cells were grown either with or without DOX for 72h. *n*≥6 biologically independent experiments. Two-sided Mann-Whitney test. **d)** Viable cell count of TC-71 and SK-N-MC cells containing a DOX-inducible re-expression construct for *TCF7L1* 72h after treatment with or without DOX. Data are mean and SEM, *n*=7 biologically independent experiments. Two-sided Mann-Whitney test. **e)** Relative percentage of area covered by colonies grown in soft-agar of TC-71 and SK-N-MC cells containing a DOX-inducible re-expression construct for *TCF7L1*. Cells were grown either with or without DOX. *n*=3 biologically independent experiments. Two-sided Mann-Whitney test. **f)** Growth of EwS subcutaneous xenografts of TC-71 and SK-N-MC cells containing a DOX-inducible re-expression construct for *TCF7L1* (arrow indicates start of DOX-treatment). Data are represented as means (*n*=7 animals/group). Two-sided Mann-Whitney test. **g)** *Ex vivo* analysis of relative necrotic area (top) and mitotic index (bottom) of xenografted TC-71 and SK-N-MC cell lines. Data are mean and SEM, *n*=7 animals/group. **h)** *Ex vivo* analysis of Ki67 positivity of xenografted TC-71 and SK-N-MC cell lines. Horizontal bars represent means and whiskers SEM, *n*=7 animals/group. *P*-values were determined via χ^2^ test testing all positives (high and moderate immunoreactivity) versus negatives. Histological images depict representative Ki67 micrographs. Scale bar=30μm.

To obtain first clues on its potential function in primary EwS, we performed Weighted Gene Correlation Network Analysis (WGCNA) based on enriched gene-sets^31^ in *TCF7L1-*correlated genes in 166 EwS tumors. Strikingly, this analysis showed that EwS tumors with low *TCF7L1* expression were enriched in embryonic pathways (**Fig. 2b**). Although *TCF7L1* is generally highly expressed in EwS, these data suggested that suppression of its transcription by EWSR1-FLI1 is associated with embryonic processes that may be linked with poor patient outcome. To test this possibility, we generated two EwS cell lines (SK-N-MC, TC-71) with a DOX-inducible re-expression of *TCF7L1*. These cell lines were chosen for functional analyses since they exhibited the lowest expression of *TCF7L1* across the 5 EwS cell lines tested (**Supplementary Fig. 1a**). As shown in **Fig. 2c**, addition of DOX to the culture medium induced a 5.5–12.8-fold re-expression of *TCF7L1*. Consistent with the hypothesis that *TCF7L1* may be a repressed EWSR1-ETS downstream transcription factor, transcriptome profiling of two EwS cell lines after either knockdown of EWSR1-ETS or re-expression of *TCF7L1* showed a highly significant (*P*=2.57×10^−165^ or *P*=5.34×10^−272^) overlap of concordantly DEGs (**Supplementary Fig. 1c**).

In agreement with this notion, conditional re-expression of *TCF7L1* was accompanied by a significant reduction of cell proliferation in both cell lines (**Fig. 2d**). Such an effect was not observed in both cell lines transduced with a DOX-inducible empty vector (**Supplementary Fig. 2a**). Moreover, long-term re-expression of *TCF7L1* significantly inhibited the capacity for clonogenic growth in two-dimensional (2D) cultures, but not in control cells (**Supplementary Figs. 2b,c**). Since stemness is a key feature enabling anchorage-independent and three-dimensional (3D) growth^32,33^, we tested whether *TCF7L1* re-expression affects spheroidal growth of EwS cells when seeded in soft-agar. As displayed in **Fig. 2e**, re-expression of *TCF7L1* significantly impaired anchorage-independent 3D growth of both EwS cell lines –an effect not observed in empty control cells (**Supplementary Fig. 2d**). Similar to these *in vitro* assays, conditional re-expression of *TCF7L1* impaired local tumor growth in a pre-clinical xenotransplantation mouse model using subcutaneous injections of tumor cells (**Fig. 2f**). Immunohistochemical (IHC) evaluation of the extracted tumors using an antibody against TCF7L1 showed specific re-expression of the protein in the DOX (+) group (**Supplementary Fig. 2e,f**). While these xenografts showed no difference in tumor necrosis in histological sections, we found a reduced mitotic index in tumors re-expressing *TCF7L1* (**Fig. 2g**). Accordingly, tumors with high *TCF7L1* expression showed a significant decrease of the proliferation marker Ki67 (**Fig. 2h**). Similar experiments performed with both cell lines transfected with empty control cells exhibited no significant differences in tumor growth, percentage of necrosis or mitotic index (**Supplementary Figs. 2g–i**).

Taken together, these results indicated that EWSR1-ETS-meditated downregulation of *TCF7L1* contributed to clonogenic and anchorage-independent growth as well as tumorgenicity of EwS cells.

### High TCF7L1 expression inhibits metastasis in EwS

Since stemness features, such as elevated clonogenic capacity and anchorage-independent growth, are essential for circulating tumor cells to colonize and to develop into clinically apparent metastases in distant organs, we reasoned that *TCF7L1* may be linked to the metastatic process in EwS. In line with this notion, transcriptome profiling, subsequent gene-set enrichment and WGCNA of SK-N-MC and TC-71 EwS cells with/without conditional re-expression of this gene uncovered that low *TCF7L1* levels led to overrepresentation of gene-sets involved in cellular migration (**Fig. 3a**). To validate this prediction, we first carried out transwell migration assays with our conditional *TCF7L1* re-expression models. Indeed, as shown in **Fig. 3b**, re-expression of *TCF7L1* significantly reduced cell migration through a porous membrane *in vitro*. In accordance, *TCF7L1* re-expression significantly inhibited invasion and single-cell 3D migration of both cell lines in fibrin gel using an advanced microfluidic chamber (**Fig. 3c**). Next, we employed an orthotopic spontaneous *in vivo* metastasis model, in which we injected SK-N-MC EwS cells with/without conditional re-expression of *TCF7L1* into the proximal tibiae of immunocompromised NOD/Scid/gamma (NSG) mice. Once signs of limping on the injected legs were observed, mice were sacrificed and legs as well as inner organs were histologically processed and evaluated for local tumor growth and evidence of macro- and micrometastasis. Excitingly, while there was no significant difference in local tumor growth in the limited space of the tibial plateau (**Supplementary Fig. 3a,b**), we observed a remarkable inhibition of macrometastatic spread to liver, lungs, and kidneys upon re-expression of *TCF7L1* (**Fig. 3d**). In fact, while 73% of organs across mice in the DOX (–) group showed evidence of metastasis to these organs, the percentage of organs with macrometastases dropped to 33% upon DOX-induced *TCF7L1* re-expression (**Fig. 3d**). In addition, we noted a significantly reduced micrometastatic burden in the same organs upon histological evaluation (**Fig. 3e**). Similar results were obtained with TC-71 cells (**Supplementary Fig. 3c–e**).

**Fig 3.**
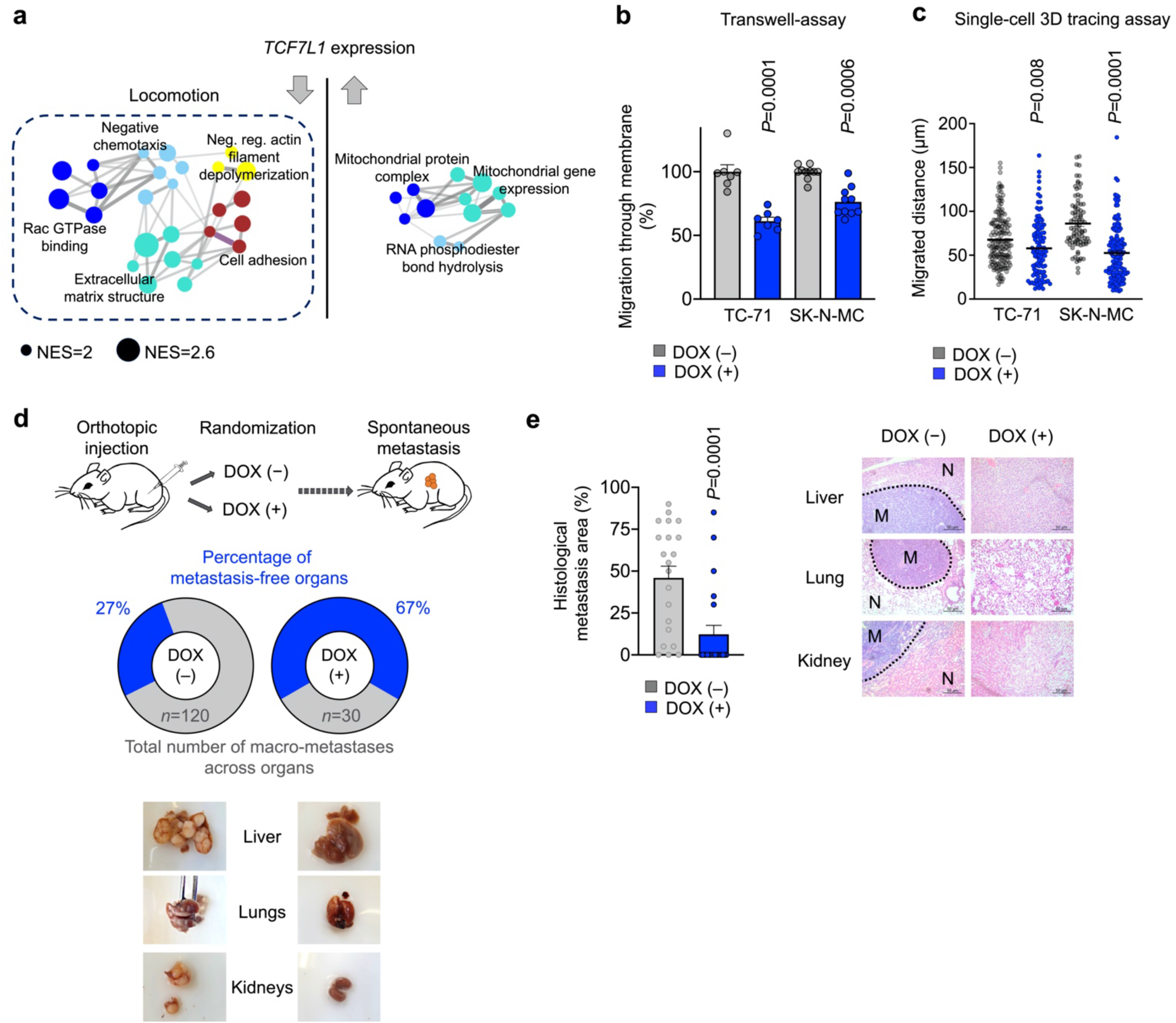
High TCF7L1 expression inhibits metastasis in EwS. **a)** Weighted Gene Correlation Network Analysis (WGCNA) depicting functional gene enrichment of down-or up-regulated genes in *TCF7L1* re-expressing EwS cells. Network depicts signatures presenting *P*<0,05, NES>2. NES, normalized enrichment score. Arrows depict direction of gene regulation. **b)** Relative percentage of migrated cells in 6h. TC-71 and SK-N-MC EwS cells containing a DOX-inducible re-expression construct for *TCF7L1* where pre-treated with or without DOX for 72h. *n*≥7 biologically independent experiments. Two-sided Mann-Whitney test. **c)** Invasion and single cell migrated distance in 15h. TC-71 and SK-N-MC EwS cells containing a DOX-inducible re-expression construct for *TCF7L1* where pre-treated with or without DOX for 72h and added to a microfluidic chamber containing a fibrin compartment. *n*≥3 biologically independent experiments. Two-sided unpaired t-test. **d)** Schematic representation of the experimental design: TC-71 or SK-N-MC EwS cell lines containing a DOX-inducible re-expression construct for *TCF7L1* were injected in the right tibia plateau. Animals were subsequently randomized and treated with or without DOX. At the end of the experiment, mice were evaluated *ex vivo* for presence of spontaneous metastases in inner organs. Pie charts depict percentage of metastasis-free organs (blue) in each condition, *n* represents total number of metastasis. Bottom pictures show representative images of metastatic organs in DOX (–) and DOX (+) conditions. *n*=8 animals/group. **e)** Graph depicts relative area of histological metastatic spread of orthotopically injected SK-N-MC EwS cells containing a DOX-inducible re-expression construct for *TCF7L1. n*=21 slides/group. Two-sided Mann Whitney test. Pictures show representative histological images of HE stainings from the evaluated organs. 20× magnification, scale bar is 50μm. M, metastasis; N, normal tissue.

In sum, these *in vitro* and *in vivo* results highlight *TCF7L1* as an inhibitor of stemness and metastasis features in EwS.

### TCF7L1 role in EwS is mediated by its DNA binding domain

To further investigate the mechanism of action of *TCF7L1*, we generated two EwS cell lines with conditional re-expression of different *TCF7L1* deletion mutants for each of its major domains (high mobility group, HMG, which is the DNA binding domain; and β-catenin binding domain, CTNNB) and subjected them to functional assays. In a first step, we tested both mutants for their phenotype in clonogenic growth assays. As depicted in **Fig. 4a**, mutants harboring a deletion of the DNA-binding domain (ΔHMG) exhibited a normal clonogenic growth capacity of TC-71 and SK-N-MC cells. However, deletion of the CTNNB domain did not show such effect (**Fig. 4a**). These data suggested that the HMG domain is required for the inhibitory effect of TCF7L1 regarding clonogenic growth. Thus, in a second step, we tested the effect of the HMG domain for the migratory and tumorigenic phenotype of both cell lines. Strikingly, in both cell lines, deletion of the DNA binding domain completely abrogated the inhibitory effect of TCF7L1 on migration and tumor growth *in vitro* and *in vivo*, respectively (**Figs. 4b**,**c**). Taken together, these data demonstrate that DNA-binding of TCF7L1 is required to suppress the tumorigenic and migratory phenotype of EwS cells.

**Fig 4.**
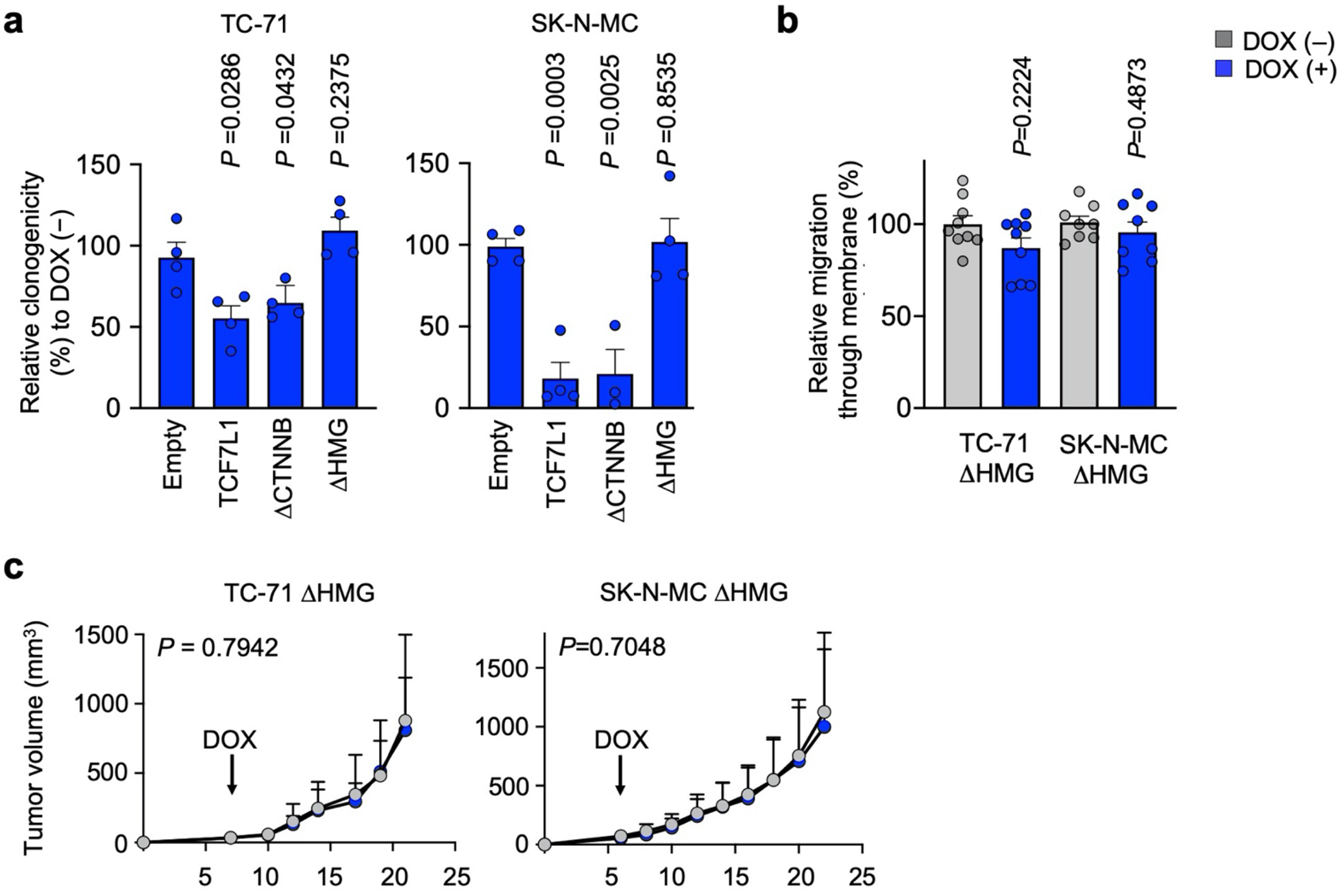
TCF7L1 role in EwS is mediated by its DNA binding domain. **a)** Relative colony number of colony-forming assays (CFAs) of TC-71 (left) and SK-N-MC (right) cells containing a DOX-inducible re-expression construct for *TCF7L1*, empty control, or one of the two deletion mutants for *TCF7L1* (deletion mutant for the β-catenin binding domain, ΔCTNNB; deletion mutant for the DNA binding domain, ΔHMG). Cells were grown either with or without DOX. *n*=4 biologically independent experiments. **b)** Relative percentage of migrated cells in 6h. TC-71 and SK-N-MC EwS cells containing a DOX-inducible re-expression construct for *TCF7L1* DNA binding domain (ΔHMG) where pre-treated with or without DOX for 72h. *n*≥8 biologically independent experiments. **c)** Growth of EwS subcutaneous xenografts of TC-71 and SK-N-MC cells containing a DOX-inducible re-expression construct for *TCF7L1* deletion mutant ΔHMG (arrow indicates start of DOX-treatment). Data are represented as means (*n*=7 animals/group). Two-sided Mann-Whitney test.

## DISCUSSION

In the present study, we harnessed a systems-biology approach to discover TCF7L1 as a key inhibitor of stemness and metastasis in EwS that is transcriptionally suppressed by EWSR1-ETS, which sheds new light on the underlying mechanisms of metastasis in this highly aggressive malignancy. Traditionally, *TCF7L1* has been described to play controversial roles in stemness/cell differentiation: on the one hand it can inhibit embryonic stem cell self-renewal through repression of pluripotency TFs^34,35^; while on the other hand, *TCF7L1* can limit the differentiation of cutaneous stem cells^36^. In cancer, the role of *TCF7L1* remains poorly understood, and appears to be cell-type specific. For example, TCF7L1 expression has been linked to promotion of cell proliferation and tumor growth and an increase of sphere formation in breast cancer^25^, colorectal cancer^26^ and skin squamous cell carcinoma^28^. In contrast, it has been shown that ectopic expression of *TCF7L1* inhibits self-renewal of liver cancer stem cells^24^, which are believed to play a prominent role in tumor relapse and metastasis^37^, thus highlighting a potential role of *TCF7L1* in the stemness-metastasis axis that strongly depends on the cellular context. Interestingly, though TCF7L1 has been described as a Wnt signaling TF^21^, in our models the CCNB1-binding domain of TCF7L1 proved to be dispensable for its inhibitory effects on clonogenic growth and migratory capacities of EwS cells, suggesting that the mechanisms that govern TCF7L1 signaling are more complex than previously anticipated. Our comparative transcriptome analyses of EwS primary tumors and TCF7L1 re-expression cell line models both highlighted an involvement of this protein in regulation of genes that collectively appear to suppress the locomotive and metastatic behavior of EwS cells. In line with this finding, low TCF7L1 expression was associated with poor patient outcome. This is particularly interesting in light of the recently proposed concept that undulating expression levels of EWSR1-FLI1 lead to a gradual switch from a more sessile and proliferative toward a more motile but less proliferative phenotype upon (transient) downregulation of the fusion oncogene^38,39^. In contrast to this binary model of EWSR1-FLI1^-high^ promoting proliferation and EWSR1-FLI1^-low^ promoting metastasis, which – to the best of our knowledge – has so far only been based on non-orthotopic, pre-clinical and cell culture-based models and has not been proven clinically, our findings support a more complex regulation of stemness and metastasis in EwS. While the previous model follows a traditional epithelial-mesenchymal-transition (EMT) concept, our findings are rather in support of a more integrative concept of the ‘migratory stem cell’^19^. In fact, our model suggests that EwS cells may reside in a ‘metastable’ state in which all EwS cells of a given EwS tumor are equipped simultaneously with both proliferative and migratory capacities that are not mutually exclusive^18^, but which may underlie a certain degree of accentuation upon different intrinsic or extrinsic cues of which EWSR1-FLI1 may be one, but not the only important component. In line with this, the notion of a clearly defined subpopulation of cancer stem cells existing universally in all cancer types has been debated in favor of a plasticity model, where more fluid, bidirectional transitions are possible. These phenotypic reversals of cancer cells at different stages of cancer progression or under different contexts has been observed in several solid tumors including melanoma, non-small cell lung cancer and glioma^40–43^. Hence, a broader view on the underlying biological mechanism of metastasis is required to fully capture this complex process in EwS. In addition, our study highlights the power of a systems biology approach to identify critical genes that would have escaped detectability by sequencing approaches, and may serve as a blueprint for similar applications in other oligomutated (pediatric) cancers.

## MATERIAL AND METHODS

### Provenience of cell lines and cell culture conditions

Human EwS cell lines and other cell lines were provided by the following repositories and/or sources: SK-N-MC, TC-71, MHH-ES1, and RD-ES cells were provided by the German Collection of Microorganism and Cell Cultures (DSMZ). A673 and HEK293T were purchased from American Type Culture Collection (ATCC). TC106 cells were kindly provided by the Children’s Oncology Group (COG). A673/TR/shEF1 cells were kindly provided by J. Alonso (Madrid)44.All cell lines were cultured in RPMI 1640 medium with stable glutamine (Biochrom, Germany) supplemented with 10% tetracycline-free fetal bovine serum (Sigma-Aldrich, Germany), 100 U/ml penicillin and 100 µg/ml streptomycin (Merck, Germany) at 37 °C with 5% CO2 in a humidified atmosphere. Cell lines were routinely tested for mycoplasma contamination by nested PCR, and cell line identity was regularly verified by STR-profiling.

### RNA extraction, reverse transcription, and quantitative real-time polymerase chain reaction (qRT-PCR)

Total RNA was isolated using the NucleoSpin RNA kit (Macherey-Nagel, Germany). 1 µg of total RNA was reverse-transcribed using High-Capacity cDNA Reverse Transcription Kit (Applied Biosystems, USA). qRT-PCR reactions were performed using SYBR green Mastermix (Applied Biosystems) mixed with diluted cDNA (1:10) and 0.5 µM forward and reverse primer (total reaction volume 15 µl) on a BioRad CFX Connect instrument and analyzed using BioRad CFX Manager 3.1 software. Gene expression values were calculated using the 2^−(ΔΔCt)^ method^45^ relative to the housekeeping gene *RPLP0* as internal control. The thermal conditions for qRT-PCR were as follows: heat activation at 95 °C for 2 min, DNA denaturation at 95 °C for 10 sec, and annealing and elongation at 60 °C for 20 sec (50 cycles), final denaturation at 95 °C for 30 sec. Oligonucleotides were purchased from MWG Eurofins Genomics (Germany) and are listed here:

*RPLP0* forward: 5’-GAAACTCTGCATTCTCGCTTC-3’

*RPLP0* reverse: 5’-GGTGTAATCCGTCTCCACAG-3’

*TCF7L1* forward: 5’-ATGAACGCCTCGATGTCC-3’

*TCF7L1* reverse: 5’-GGTTCCTGCTTGACGATGG-3’

*EWSR1-FLI1* forward: 5’-GCCAAGCTCCAAGTCAATATAGC-3’

*EWSR1-FLI1* reverse: 5’-GAGGCCAGAATTCATGTTATTGC-3’

### *TCF7L1* overexpression experiments

To re-express *TCF7L1* at physiological levels, we assessed the baseline expression levels of *TCF7L1* in 18 EwS cell lines for which whole-transcriptome data from Affymetrix Clariom D arrays was available (triplicates per cell line). As suitable models, we chose SK-N-MC and TC-71 cells as they exhibited the lowest baseline *TCF7L1* expression among these cell lines and proceeded with cloning as described in^46^. Briefly, full-length cDNA of *TCF7L1* (NM_031283) was PCR-amplified from a commercial plasmid (Origene, SC126274) and an HA-tag was added by using AgeI-and NotI-restriction site containing primers (forward: 5’-ATTAACCGGTGCCACCATGCCCCAGCTCG-3’; revers: 5’-TAATGCGGCCGCTTAAGCGTAATCTGGAACATCGTAGTGGGCAGACTTGGTGACC −3’) and a final Tm of 65 °C (Phusion Polymerase, ThermoFisher), before cloning it into the multiple cloning site of a modified pTP vector^47^. *TCF7L1* non-functional mutants were cloned from the same *TCF7L1* (NM_031283) ORF cDNA clone using touchdown-PCR. The HA-tagged CTNNB1-binding site truncated mutant was generated by using a different forward primer (5’-ATTAACCGGTGCCACCATGAACCAGAGCAGCAGCT-3’) and a final Tm of 63 °C. The HMG-Box binding site was deleted by fusing two PCR products (Product 1, final Tm 57 °C): forward: 5’-ATTAACCGGTGCCACCATGCCCCAGCTCG-3’; reverse: 5’-CTTACCATAGTTGTCGGGCTTCTTTTCCTCCT-3’; Product 2 (final Tm 64 °C): forward: 5’-GAGGAAAAGAAGCCCGACAACTATGGTAAGAAAAAGAAGAGGA-3’; reverse: 5’-TAATGCGGCCGCTTAAGCGTAATCTGGAACATCGTAGTGGGCAGACTTGGTGACC −3’) using an overhang-extension PCR protocol (overhang-extension PCR (final Tm 65 °C): forward: 5’-ATTAACCGGTGCCACCATGCCCCAGCTCG-3’; reverse: 5’-TAATGCGGCCGCTTAAGCGTAATCTGGAACATCGTAGTGGGCAGACTTGGTGACC −3’). The generated inserts were double restriction-digested with AgeI and NotI (NEB) and ligated into the pTP backbone^47^ using T4 ligase (NEB). Positive clones were identified by colony PCR and cultured in 100 ml of LB Broth containing 100 µg/ml Ampicillin. Plasmids were extracted and purified using a Midi-Prep Kit (Macherey-Nagel). The correct insertion of full-length *TCF7L1* cDNA, empty control or *TCF7L1*-deficient mutants was verified by Sanger sequencing (sequencing primers: forward 5’-ACGTATGTCGAGGTAGGCGT-3’; reverse 5’-TTCGTCTGACGTGGCAGC-3’). Lentiviral particles were generated in HEK293T cells and used for transduction of SK-N-MC and TC-71 EwS cells using polybrene (8 µg/ml). Transduced cells were selected with 0.5 µg/ml puromycin. Re-expression of *TCF7L1* in SK-N-MC and TC-71 cells was achieved by addition of 1 µg/ml DOX to the culture medium. Cells were single-cell cloned and specific clones were selected and tested for re-expression of *TCF7L1*. Only the clones exhibiting re-expression levels similar to the expression FCs after *EWSR1-FLI1* silencing (see above) were selected for functional assays. Cells transfected with the empty vector were used as additional controls.

### Transcriptome analyses

To assess the potential effect of *TCF7L1* on gene expression in EwS cells, microarray analysis was performed. To this end, 1.2×10^4^ cells per well were seeded in 6-well plates and treated with 1μg/μl DOX for 72h (DOX-refreshment after 48h). Thereafter, total RNA was extracted with the ReliaPrep miRNA Cell and Tissue Miniprep System (Promega) and RNA quality was assessed with a Bioanalyzer. All samples had an RNA integrity number (RIN)>9 and were hybridized to Human Affymetrix Clariom D microarrays. Gene expression data were quantile normalized with Transcriptome Analysis Console (v4.0; Thermo Fisher Scientific) using the SST-RMA algorithm as previously described^48^. Annotation of the data was performed using the Affymetrix library for Clariom D Array (version 2, Homo sapiens) on gene level. DEGs with consistent and significant FCs across cell lines were identified as follows: Normalized gene expression signals were log2 transformed. To avoid false discovery artifacts due to the detection of only minimally expressed genes, we excluded all genes with a lower expression value than that observed for *ERG* of the respective cell lines (log2 expression signal of 7.08 for SK-N-MC and 6.57 for TC-71), which is known to be virtually not expressed in EWSR1-FLI1 positive EwS cell lines^15^. The FCs of the control samples (empty vector) and both *TCF7L1* re-expressing EwS cell lines were calculated for each cell line separately. Then the FCs in the *TCF7L1* re-expressing samples were normalized to that of the empty control cells. Then both FCs were averaged to obtain the mean FC per gene across cell lines. DEGs were determined as having a log2 FC >0.5 or <–0.5, respectively.

### Gene-set enrichment analysis (GSEA)

To identify enriched gene-sets, genes were ranked by their expression FC between the groups DOX (−) and DOX (+). GSEA was performed using the FGSEA R package (v 3.6.3) based on Gene Ontology (GO) biological processes terms from MSigDB (c5.all.v7.0.symbols.gmt)^49^. GO terms were filtered for statistical significance (adjusted *P*<0.05) and a normalized enrichment score |(NES)|>2 (10,000 permutations). In order to construct a network, the Weighted Gene Correlation Network Analysis R package (WGCNA R)^31^ was used. Briefly, a binary matrix of GO-terms × genes (where 1 indicates the gene is present in the GO term and 0 indicates it is not) was created. Then, the Jaccard’s distance for all possible pairs was computed to create a symmetric GO adjacent matrix. Clusters of similar GO terms were identified using dynamicTreeCut algorithm, and the top 20% highest edges were selected for visualization. The highest scoring node in each cluster was determined as the cluster label (rName). The obtained network and nodes files were fed into Cytoscape (v 3.8.0) for network design and visualization as previously described^50^.

### Correlation analysis

To identify gene-sets correlated with *TCF7L1* expression in primary tumors, we created a pre-ranked list of genes ordered by Pearson’s correlation coefficient with *TCF7L1* based on transcriptomic data of 166 of EwS^23^ and GSEA was performed as described above.

### Proliferation assays

For proliferation assays, 5–8×10^5^ EwS cells per well (depending on the cell line) were seeded in triplicates per group in 6-well plates and treated with 1 µg/ml DOX for 72h. Thereafter, cells including their supernatant were harvested and counted using standardized hemocytometers (C-Chip, Biochrom) and the Trypan-Blue (Sigma-Aldrich) exclusion method as described in^51^.

### Clonogenic growth assays

For clonogenic growth assays, *TCF7L1* re-expressing EwS cells and respective controls were seeded in triplicates at low density (2×10^3^ cells) per well in 12-well plates and grown for 9–11 days (depending on the cell line) with/without DOX-treatment (renewal of DOX or vehicle every 48h). Thereafter, colonies were stained with crystal violet (Sigma-Aldrich) and colony numbers and areas were measured with the ImageJ Plugin *Colony area*. The clonogenicity index was calculated by multiplying the counted colonies with the corresponding colony area.

### Sphere formation assays in soft agar

For the analysis of anchorage-independent growth, *TCF7L1* re-expressing EwS cells and respective controls were pre-treated with/without DOX for 48h before seeding. A base of 2 ml of agar 1:1 with 2× DMEM medium was poured in wells of 6-well plates and left for 30–60 min to solidify. Then, 5×10^3^ cells per well were seeded in 500 µl in triplicates per condition (– /+DOX). Cells were kept in culture for 9–11 days and new medium (–/+DOX) was added on top of each well every 48h. Spheres were stained with 50µl of a 5 mg/ml MTT solution that was added dropwise to each well and incubated for 1h. Pictures of the stained spheres were taken and their area was analyzed with ImageJ.

### Transwell assays

For the analysis of migration through a porous membrane we proceeded as described in^52^. Briefly, *TCF7L1* re-expressing EwS cells, their mutants, and respective controls were pre-treated with/without DOX for 72h before seeding and were starved by decreasing FCS concentration in the growth medium to 0.5% during the last 24h. The following morning, fresh medium with 10% FCS (chemoattractant) was plated in the lower compartment of a 24-well plate. Transwell attachments were placed on top and 1×10^5^ cells were plated on them with medium containing 0.5% FCS with/without DOX. Cells were allowed to migrate for 6h. After that, transwell inserts were washed with PBS and the top membrane was cleaned of remaining unmigrated cells with a cotton swab. Migrated cells were fixed with a 4% formaldehyde solution and stained with crystal violet. Transwells were washed with water to eliminate excess crystal violet and migrated cells were de-stained with a solution containing 2% acetic acid in PBS during 10 minutes on a shaker. The obtained solution was measured by absorbance at 570 nm in a plate reader.

### Microfluidic invasion and single cell 3D migration assay

TC-71 and SK-N-MC *TCF7L1* re-expressing EwS cells were pre-treated for 48h with DOX (1µg/ml). Then, 1.5×10^6^ cells/ml were embedded in a 2.5 mg/ml fibrin matrix and introduced in the central channel of a microfluidic device as described^53^. Following fibrin polymerization (20 min inside a humidity chamber), culture medium was added to the side channels of the microfluidic device. The next day, medium was substituted with CO2 independent culture medium (Thermo Fisher) and cells were imaged with an automated wide-field epifluorescence microscope (Lionheart FX, BioTek Instruments, Inc., Winooski, VT, USA). Images of at least 5 regions of interest for each microfluidic device per condition (*n*=3) were captured every 30 min for 15h. Temporal stacks were reconstructed using ImageJ and single cell 3D movement inside the fibrin matrix was quantified employing the image processing software Imaris (v.9.1, Bitplane AG, Zurich, Switzerland).

### *In vivo* experiments in mice

3×10^6^ SK-N-MC or TC-71 EwS cells harboring a re-expression construct for either *TCF7L1*, the *TCF7L1* deletion mutant (*TCF7L1*-HMGmut) or empty controls were injected in a 1:1 mix of cells suspended in PBS with Geltrex Basement Membrane Mix (ThermoFisher) in the right flank of 10–12 weeks old NOD/scid/gamma (NSG) mice as described in^54^. Tumor diameters were measured every second day with a caliper and tumor volume was calculated by the formula L×l^2^/2, where L is the length and l the width. When the tumors reached an average volume of 80 mm^3^, mice were randomized in two groups of which one was henceforth treated with 2 mg/ml DOX (Beladox, Bela-pharm, Germany) dissolved in drinking water containing 5% sucrose (Sigma-Aldrich) to induce an *in vivo* re-expression (DOX (+)), whereas the other group only received 5% sucrose (control, DOX (−)). Once tumors of control groups reached an average volume of 1,500 mm^3^, all mice of the experiment were sacrificed by cervical dislocation. Other humane endpoints were determined as follows: Ulcerated tumors, loss of 20% body weight, constant curved or crouched body posture, bloody diarrhea or rectal prolapse, abnormal breathing, severe dehydration, visible abdominal distention, obese Body Condition Scores (BCS), apathy, and self-isolation. For analysis of EwS growth in bone, EwS cells were orthotopically injected into the proximal tibial plateau of NSG mice. To this end, one day before injection, mice were pre-treated with 800 mg/kg mouse weight/day Metamizol in drinking water as analgesia. On the day of injection, mice were anesthetized with inhaled isoflurane (1.5-2.5% in volume) and their eyes were protected with Bepanthen eye cream. After disinfection of the injection site, 2×10^5^ cells/20µl were directly injected with a fine 28 G needle (Hamilton, USA) into the right proximal tibia. For durable pain prophylaxis until the first day after intraosseous injection, mice were subsequently treated with Metamizole in drinking water (800 mg/kg mouse weight/day). At the first day after injection of tumor cells, mice were randomized in two groups of which one received henceforth 2 mg/ml DOX (BelaDox, Bela-pharm) dissolved in drinking water containing 5% sucrose (Sigma-Aldrich) to induce *TCF7L1* re-expression (DOX (+)), whereas the other group only received 5% sucrose (control, DOX (−)). All tumor-bearing mice were sacrificed by cervical dislocation at the predefined experimental endpoint, when the mice reached a humane endpoint as listed above or exhibited signs of limping at the injected leg (event). After extraction of the tumors, a small fraction of each tumor was snap frozen in liquid nitrogen to preserve the RNA isolation, while the remaining tumor tissue was fixed in 4%-formalin and embedded in paraffin for immunohistology. In the case of the orthotopic model, all inner organs were harvested, weighed, photographed, 4%-formalin-fixed and embedded in paraffin for (immuno)histology. For analysis of the extent of metastatic spread, HE-stained histological slides were evaluated for presence of EwS cells and the area of metastasis versus total area was calculated. Animal experiments were approved by the governments of Upper Bavaria and Northbaden and conducted in accordance with ARRIVE guidelines, recommendations of the European Community (86/609/EEC), and United Kingdom Coordinating Committee on Cancer Research (UKCCCR) guidelines for the welfare and use of animals in cancer research.

### Immunohistochemistry (IHC) and immunoreactivity scoring (IRS)

For IHC, 4-µm sections were cut and antigen retrieval was carried out by heat treatment with Target Retrieval Solution (S1699, Agilent Technologies) for Ki67, or Tris buffer pH 9.0 for TCF7L1 staining, respectively. For Ki67, the slides were stained with monoclonal anti-Ki67 raised in rabbit (1:200, 275R-15, Cell Marque/Sigma-Aldrich) for 60 min at RT, followed by a monoclonal secondary horseradish peroxidase (HRP)-coupled horse-anti-rabbit antibody (ImmPRESS Reagent Kit, MP-7401, Vector Laboratories). AEC-Plus (K3469, Agilent Technologies) was used as chromogen. For TCF7L1, the slides were stained with polyclonal anti-TCF7L1 antibody raised in rabbit (1:100 for complete slides, 1:25 for TMAs; 28835 clone D15G11 lot 5, Cell Signaling). For TMA analysis, DAKO REAL Detection System, Alkaline Phosphatase/RED Rabbit/Mouse was used (K5005, Dako). For xenograft slides, Biotin-SP-conjugated AffiniPure Goat Anti-Rabbit IgG (H+L) (111-065-144, Jackson Immunoresearch) and Alkaline Phosphatase Streptavidin (SA-5100, Vector) were used. The chromogen used was that included in the DAKO REAL Detection System (K 5005, Dako). Samples were counterstained with hematoxylin (H-3401, Vector Laboratories; or T865.3, Mayer, Roth). FFPE xenografts of the EwS cell lines were stained with HE for mitosis counting and metastatic spread assessment. Slides were scanned on a Nanozoomer-SQ Digital Slide Scanner (Hamamatsu Photonics K.K.) and visualized using NDP.view2 image viewing software (Hamamatsu Photonics K.K.). Mitoses were quantified by a blinded observer per high-power field (40×). Final scores/quantifications were determined by examination of 5–10 high-power fields of at least one section per sample. Evaluation of immunoreactivity of TCF7L1 was carried out in analogy to scoring of hormone receptors with a modified Immune Reactive Score (IRS) ranging from 0–12 that was adapted to EwS tumors^55,56^. Briefly, the percentage of cells with expression of the given antigen was scored and classified in five grades (grade 0=0−19%, grade 1=20−39%, grade 2=40−59%, grade 3=60−79% and grade 4=80−100%). In addition, the intensity of marker immunoreactivity was determined (grade 0=none, grade 1=low, grade 2=moderate and grade 3=strong). The product of these two grades defined the final IRS. Slides were scored by *n*≥2 independent observers.

### Survival analysis

Kaplan-Meier survival analyses were carried out in 166 EwS patients whose molecularly confirmed and retrospectively collected primary tumors were profiled at the mRNA level by gene expression microarrays in previous studies^57–60^. To that end, microarray data generated on Affymetrix HG-U133Plus2.0, Affymetrix HuEx-1.0-st or Amersham/GE Healthcare CodeLink microarrays of the 166 EwS tumors (Gene Expression Omnibus (GEO) accession codes: <u>GSE63157</u>^57^, <u>GSE12102</u> ^58^, <u>GSE17618</u> ^59^, <u>GSE34620</u> ^60^ provided with clinical annotations were normalized separately as previously described^55^. Only genes that were represented on all microarray platforms were kept for further analysis. Batch effects were removed using the ComBat algorithm^61^. Data processing was done in R.

### Statistical analysis and software

Statistical data analysis was performed using PRISM 9 (GraphPad Software Inc., Ca, USA) on the raw data. If not specified otherwise in the figure legends, comparison of two groups in functional *in vitro* experiments was carried out using a two-sided Mann-Whitney test. If not specified otherwise in the figure legends, data are presented as dot plots with horizontal bars representing means and whiskers representing the standard error of the mean (SEM). Sample size for all *in vitro* experiments were chosen empirically. For *in vivo* experiments, the sample size was predetermined using power calculations with *β*=0.8 and *α*<0.05 based on preliminary data and in compliance with the 3R principles (replacement, reduction, refinement). In Kaplan-Meier survival analyses, curves were calculated from all individual survival times of mice, respectively. Statistical differences between the groups were assessed by a Mantel-Haenszel test.

## Supporting information

Supplementary Tables_Cidre-Aranaz et al.

## Data availability

Original microarray data that support the findings of this study were deposited at the National Center for Biotechnology Information (NCBI) GEO and are accessible through the accession number GSE165929.

## ACKNOWLEDGEMENTS

We thank Dr. Paul Northcott and Dr. Brian Gudenas for providing R scripts for weighted gene correlation network analyses and Carlos Rodriguez-Martin for assistance with script adaptation. We thank Anja Heier and Andrea Sendelhofert for expert technical assistance. This work was mainly supported by the Wilhelm Sander-Foundation (2016.167.1), the Barbara & Hubertus Trettner Foundation, and the Dr. Rolf M. Schwiete Foundation. In addition, the laboratory of T.G.P.G. was supported by the Matthias-Lackas Foundation, the Dr. Leopold and Carmen Ellinger Foundation, the German Cancer Aid (DKH-70112257 and DKH-70114111), the Gert und Susanna Mayer Foundation, the SMARCB1 association, and the Barbara und Wilfried Mohr Foundation. J.L. was supported by a scholarship of the Chinese Scholarship Council (CSC), and M.K. and C.M.F. by scholarships from the German Cancer Aid. M.M. was supported by the Swiss National Science Foundation (SNF 310030_179167).

## AUTHOR CONTRIBUTIONS

F.C.A, M.M.L.K., and T.G.P.G. conceived the study. F.C.A. and T.G.P.G wrote the paper, and drafted the figures and tables. F.C.A., M.M.L.K., and S. K. carried out *in vitro* experiments. F.C.A, M.F.O., and T.G.P.G. performed bioinformatic and statistical analyses. F.C.A, M.M.L.K., J.L., T.L.B.H., J.M, R.I., and A.B. performed and/or coordinated *in vivo* experiments. M.F.O., K.C., M.J.C.G., G.A., S.B., C.A., M.M., C.F., A.M., S.O., L.R.-P., and M.K. contributed to experimental procedures. T.K. and F.B. provided laboratory infrastructure and/or histological guidance. T.G.P.G. supervised the study and data analysis. All authors read and approved the final manuscript.

## LEGENDS TO SUPPLEMENTARY FIGURES

**Supplementary Fig 1.**
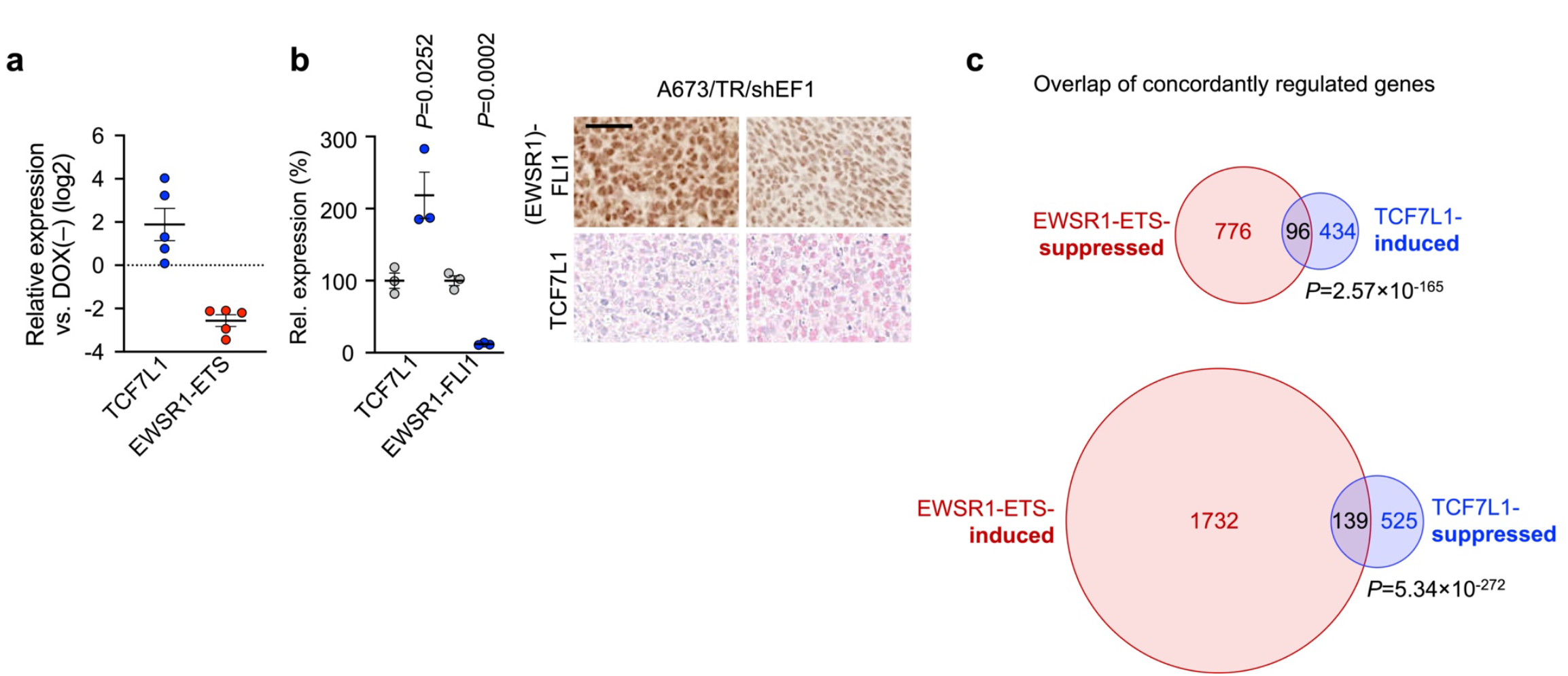
TCF7L1 expression in EwS cell lines. **a)** *TCF7L1* and *EWSR1-ETS* expression in 5 EwS cell lines with conditional knock-down (KD) of EWSR1-ETS for 72h measured by qRT-PCR. *n*≥*3* biologically independent experiments. **b)** Left: *EWSR1-FLI1* and *TCF7L1* expression (Affymetrix microarrays) in A673/TR/shEF1 xenografts after 96 h of DOX treatment. Horizontal bars represent means, *n*= 3 xenografts per group. *T*wo-sided independent *t*-test. Right: representative IHC for (EWSR1-)FLI1 and TCF7L1. Scale bar = 20 µm. **c)** Size-proportional Venn diagrams of genes concordantly regulated by 96h after knockdown of *EWSR1-ETS* or upregulation of *TCF7L1* in TC-71 and SK-N-MC EwS cells. Minimum log2 fold-change ± 0.5. Fischer’s exact test.

**Supplementary Fig 2.**
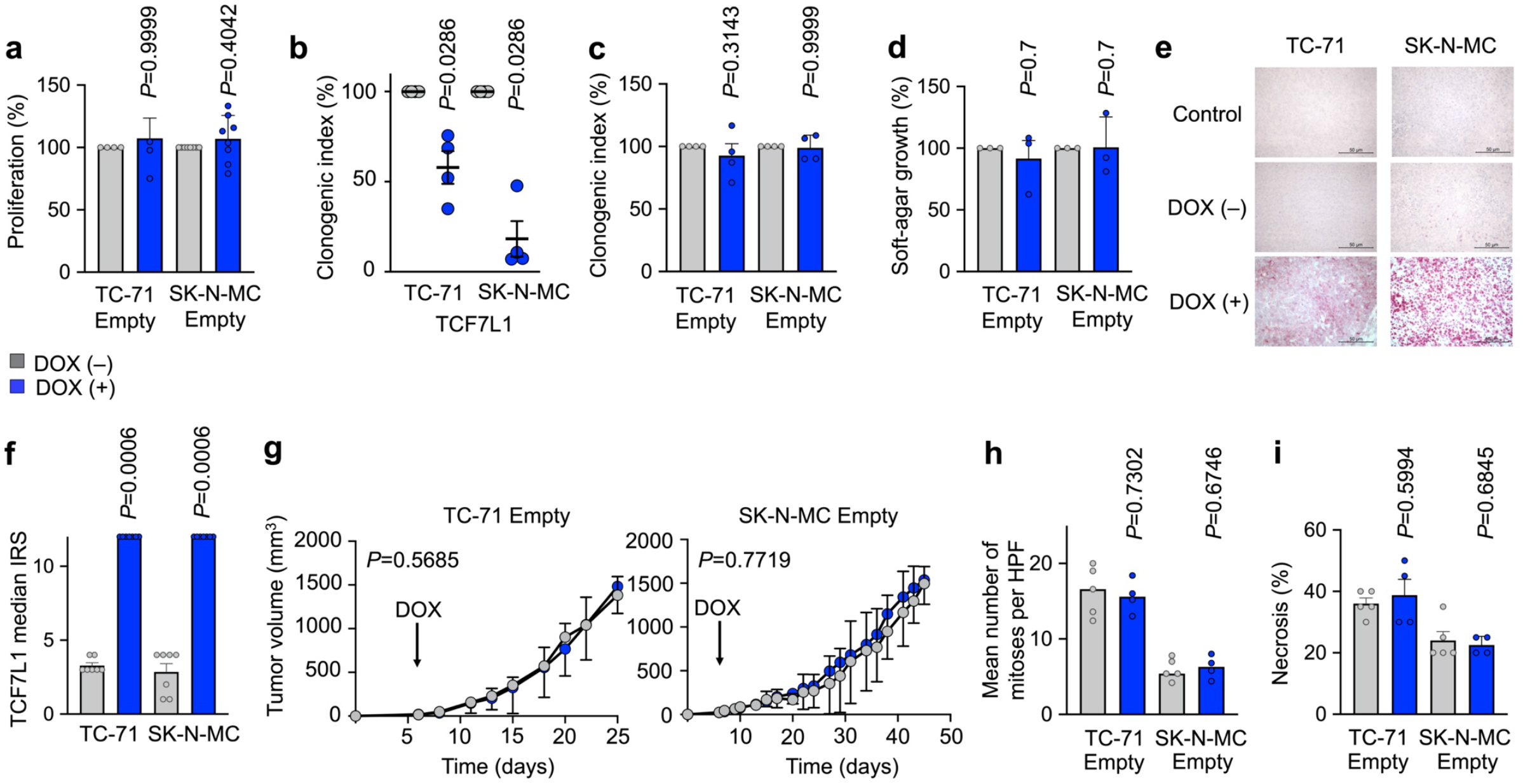
TCF7L1 re-expression inhibits tumorigenesis *in vitro* and *in vivo*. **a)** Viable cell count of TC-71 and SK-N-MC cells containing a DOX-inducible re-expression construct for an empty control 72h after treatment with or without DOX. Data are mean and SEM, *n*≥*3* biologically independent experiments. Two-sided Mann-Whitney test. **b)** Relative colony number of colony-forming assays (CFAs) of TC-71 and SK-N-MC cells containing a DOX-inducible re-expression construct for *TCF7L1*. Cells were grown either with or without DOX. *n*=4 biologically independent experiments. Two-sided Mann–Whitney test. **c)** Relative colony number of CFAs of TC-71 and SK-N-MC cells containing a DOX-inducible re-expression construct for an empty control. Cells were grown either with or without DOX. *n*=4 biologically independent experiments. Two-sided Mann-Whitney test. **d)** Relative percentage of area covered by colonies grown in soft-agar of TC-71 and SK-N-MC cells containing a DOX-inducible re-expression construct for an empty control. Cells were grown either with or without DOX. *n*=3 biologically independent experiments. Two-sided Mann-Whitney test. **e)** Representative images of *TCF7L1* expression on protein level by IHC on xenografted tissues from *in vivo* subcutaneous xenografts of TC-71 and SK-N-MC EwS cells with conditional re-expression of TCF7L1. **f)** Immuno Reactive Scores (IRS) of TCF7L1 expression in all subcutaneous xenografts of TC-71 and SK-N-MC EwS cells with conditional re-expression of TCF7L1 (*n*=7 animals/group). Horizontal bars represent mean. **g)** Subcutaneous growth of EwS xenografts of TC-71 and SK-N-MC cells containing a DOX-inducible re-expression construct for an empty control (arrow indicates start of DOX-treatment). Data are represented as means (*n*=5 animals/group). Two-sided Mann-Whitney test. **h)** *Ex vivo* analysis of mitotic index. Data are mean and SEM, *n*≥4 animals/group. **i)** Relative necrotic area of xenografted empty control TC-71 and SK-N-MC cell lines. Data are mean and SEM, *n*≥4 animals/group.

**Supplementary Fig 3.**
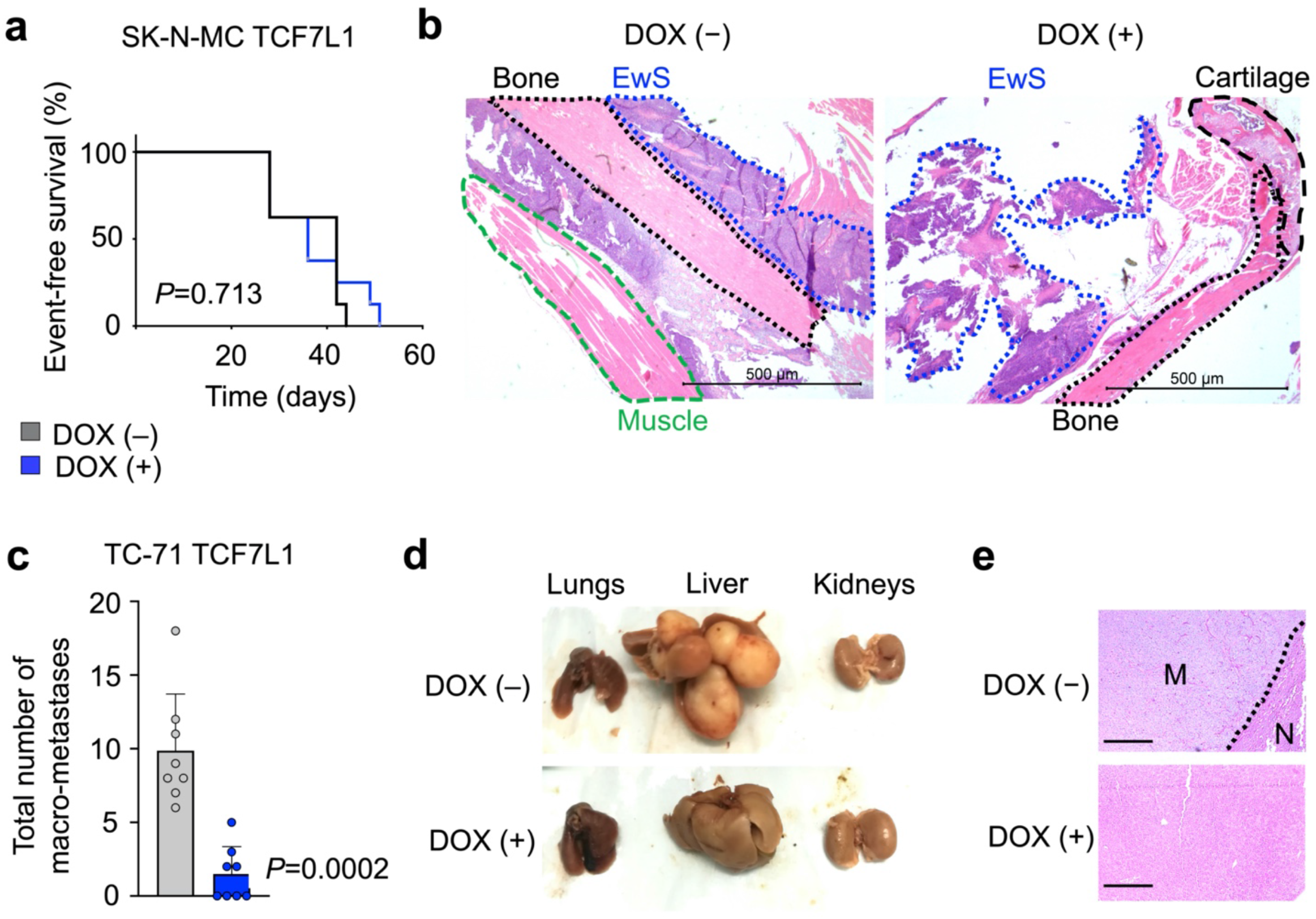
High TCF7L1 expression inhibits metastasis in EwS. **a)** Kaplan-Meier plots showing event-free survival time of NSG mice bearing orthotopic xenografts of SK-N-MC cells containing a DOX-inducible re-expression construct for *TCF7L1*. After intra-osseous injection, mice were randomized and treated with either vehicle (–) or DOX (+). Event: first signs of limping. Log-rank (Mantel-Cox) test **b)** Representative micrographs of HE from tibias obtained from both groups of mice orthotopically injected with SK-N-MC cells containing a DOX-inducible re-expression construct for *TCF7L1*. Mice were treated with either vehicle (–) or DOX (+). Scale bar=500 µm. EwS infiltrating area is shown in blue dots, black dots show bone tissue, green lines show muscle tissue and black lines show cartilage tissue. **c)** Graph depicts total number of macro-metastases counted after inner organ extraction in mice bearing orthotopically implanted TC-71 cells containing a DOX-inducible re-expression construct for *TCF7L1. n*=8 evaluated livers from independent animals in each group. **d)** Representative images from metastatic organs of mice bearing orthotopically implanted TC-71 cells containing a DOX-inducible re-expression construct for *TCF7L1* in DOX (–) and DOX (+) conditions. **e)** Representative histological images of HE stainings from the evaluated organs. 20× magnification, scale bar=50μm. M, metastasis; N, normal tissue.

